# Functionally significant, novel variants of *BMP4* are associated with isolated congenital heart disease

**DOI:** 10.1101/2025.05.26.656099

**Authors:** Jyoti Maddhesiya, Ritu Dixit, Ashok Kumar, Bhagyalaxmi Mohapatra

**Affiliations:** Cytogenetics Laboratory, Department of Zoology, Institute of Science, Banaras Hindu University, Varanasi, Uttar Pradesh, India; Department of Pediatrics, Institute of Medical Sciences, Banaras Hindu University, Varanasi, Uttar Pradesh, India

**Keywords:** Congenital heart disease, BMP4, SMAD, TGF-β, variations

## Abstract

Bone morphogenetic proteins (BMPs) are multipotent cytokines of TGFβ super family, involved in wide range of biological processes including embryonic development, tissue differentiation, cell proliferation, migration and organogenesis. BMP signaling also plays a pivotal role during different phases of cardiogenesis. Although genetic variations in several components of BMP signaling have been linked to congenital heart disease (CHD), many of these findings lack thorough functional validation. To assess the role of BMPs in CHD, Sanger sequencing of *BMP4* gene was conducted in 285 CHD cases along with 400 healthy controls. Four missense novel pathogenic variants were detected in four unrelated CHD probands with heterogeneous phenotypes. All the four variants (p.R113G, p.E151V, p.T197I and p.R226W) were located in BMP4 pro-peptide domain. Western blotting analysis revealed a significant increase in the phosphorylation of SMAD1/5 caused by all the four variants. Further, all the four variants enhance the transactivation of BMP-responsive promoters, *Id1-luc* and *Id3-luc* in luciferase reporter assay. Moreover, qRT-PCR analysis validating the enhanced endogenous expression of downstream targets namely *Smad1*, *Smad5*, *Id1*, *Id3*, and *Irx4* which further confirm the augmented activity due to all the four variants. Besides, our computational modeling of RNA structures and its features, modifications in secondary and tertiary structures and various physiochemical properties are speculated to enhanced the binding of BMP4 muteins with its respective partner and thereby boosting the SMAD-dependent BMP signaling. Altogether, both *in vitro* and *in silico* observations unveiling the gain-of-function activity of mutants which potentially perturbing the normal BMP signaling/ dynamics, consequently inducing CHD.

## 1. Introduction

Congenital heart diseases (CHDs) are a group of structural and functional defects in cardio-vascular system that occur during the development of the heart, affecting ∼1% of all live new-borns [1]. Increasing evidence underscore the role of genetic factors underlying most of cases of CHD [2–5]. During embryogenesis, the heart development undergoes an amalgamating signaling network coordinated by the transforming growth factor-β (TGFβ) superfamiy, including bone morphogenetic protein (BMP), nodal growth factor (NODAL), growth and differentiation factors (GDFs), Wnt family member, and fibroblast growth factor (FGF) and many others. all together which tightly regulate the expression of cardiac enrich transcription factors like *NKX2.5*, *GATA4/5/6* and *TBXs*, *IRX, MEFs, SRF* and *CITED2, etc.* The mutual cooperation of these promote embryonic growth, and differentiation, which include determination of heart fields, mesodermal differentiation of cardiac progenitor cells into cardiomyocytes, endocardial cushion formation leading to formation of four chambered functional heart [4,6].

Bone morphogenetic proteins (BMPs), member of TGFβ super family, are highly conserved signaling molecules, play important role during embryonic growth and development [7]. Although BMPs are known for their role in osteogenesis and bone growth, these are multipotent factors involved in diverged cellular functions during embryonic and adult life including cell proliferation, differentiation, migration, apoptosis, patterning of body axis and organogenesis. Accumulating evidences suggest crucial role of BMP signaling during heart development and cardiomyocyte differentiation [8]. Animal studies have demonstrated that heterozygous mutant mouse for *Bmp2* and *Bmp4* develop valve and septal defects [9]. Further *Bmp2* and *Bmp4* have been detected to express in endocardial cushion [10–12]. BMP signaling is involved in endocardium activation following epithelial to mesenchymal transformation (EMT), promotion and invasion of the mesenchyme into cardiac cushions [13].

The first evidence of BMP signaling in heart development was derived from the expression of *dpp* gene in dorsal ectoderm, which control the mesodermal expression of *tinman,* essential for heart tube formation in Drosophila [14]. However, the expression of *Dpp* and *BMP4* are noted to be in opposite pattern in vertebrates [15]. Chordin, a vertebrate homologue of Drosophila *sog* gene, antagonise BMP4 and restrict it in the ventral half of the embryo [16]. During embryonic heart development, *Bmp4* has been reported to express by E8.5 in myocardium of the outflow tract (OFT) and continue to express throughout the heart development and remodelling [10]. It also plays critical role in OFT and atrio-ventricular septation [17]. *BMP4* also plays an important role during the differentiation of cardiomyocyte progenitors into cardiomyocytes. Nonetheless, its effect at commitment stages is dependent on a precise balance, with other laterality genes namely activin A, Nodal, and Wnt signaling [18].

Posch et al., (2008), first associated a variation (p. V152A) in *BMP4* in patients with congenital cardiac septal defects [19]. Subsequently, several other variations have also been discovered in *BMP4* gene in patients with CHD [20–22]. Besides, *BMP4* have been associated with several other malformations namely cleftlip, eye, brain, kidney, tooth agenesis and bone defects [23–27].

In the present study, we report four novel variations identified in *BMP4* in CHD patients from Indian population. All four variations reside in the pro-domain of the protein. *In silico* analyses demonstrate remarkable structural deformities of the protein, simultaneously *in vitro* functional studies implicate a gain-of-function activity of the BMP signaling.

## 2. Materials and Methodology

### 2.1 Participant recruitment and clinical investigation

A total of 285 isolated CHD patients (111 females and 174 males with a median age of 3 years, ranging from 0-30 years of age) were enrolled including 8 familial and 277 sporadic patients along with 400 healthy unrelated age-matched control individuals (median age 3.7 years, ranging from 1 to 32 years of age) from the same geographical locality after intensive investigation about their family history and clinical diagnosis by 2D-colour doppler echocardiography, ECG, and chest X-ray. All the individuals were recruited from Department of Cardiology and the Department of Paediatric Medicine, SS Hospital, Banaras Hindu University (BHU), Varanasi, India after obtaining written informed consent. The Institutional Human Ethical Committee of the University has ethically approved the study. The recruited probands possess miscellaneous CHD phenotype which is tabulated in **Supplementary Table 1.** Probands associated with syndromes were eliminated from the study.

### 2.2 Screening, identification, and analysis of variants

The genomic DNA samples from peripheral blood leukocytes were purified by ethanol precipitation method from all the study subjects. All coding exons as well as the splicing boundaries of human *BMP4* (referential genomic DNA sequence, NC_000014.9) were PCR amplified. Following enzymatic purification of the Amplified PCR products, by Exonuclease I (USB Products, Affymetrix, Inc. USA) and recombinant Shrimp Alkaline Phosphatase (USB Products, Affymetrix, Inc.), Sanger sequencing was conducted on ABI PRISM 3130 genetic analyser (Applied Biosystems, USA) using Big Dye® Terminator v3.1 Cycle Sequencing Kit (Applied Biosystems, Inc. USA). The chromatogram peaks were analysed by Finch TV software (http://www.geospiza.com/ftvdlinfo. html, Geospiza). The detected variants were verified by a second-independent PCR and sequencing analysis. The inheritance patterns were analysed by parental genotyping. Different databases such as gnomAD (https://gnomad.broadinstitute.org/), ClinVar (http://www.ncbi.nlm. nih.gov/clinvar/) and dbSNP (https://www.ncbi.nlm.nih.gov/snp/) and 1000G (https://www. internationalgenome.org/) databases from National Centre for Biotechnology Information (NCBI) as well as INDEX-db (http://indexdb.ncbs.res.in/), GenomeAsia100K (https://browser.genomeasia100k.org/) were consulted to verify the novelty of identified *BMP4* variants.

### 2.3 Characterizing BMP4 variants by in silico analysis

The disease-causing potential of variants as well as its effects on the stability, structure, and function of proteins were evaluated by different *in silico* tools. To carry out all the *in silico* analyses, BMP4 reference genomic DNA (NC_000014.9) and mRNA sequence (NM_001202.6) from Gen-Bank (https://www.ncbi.nlm.nih.gov/genbank/) and protein sequence (NP_001193.2) from Protein database (https://www.ncbi.nlm.nih.gov/protein/) were fetched.

#### 2.3.1 Phylogenetic conservation analysis of BMP4 mutated amino acids

The conservation of amino acid altered by the missense variants were checked by searching the reference protein sequence of BMP4 from Protein database (https://www.ncbi.nlm.nih.gov/protein/). Using NCBI ‘Homologene’ feature, cross species alignment of human BMP4 protein was performed with different species of vertebrates such as *H. sapiens, P. troglodytes, M. mulatta, C. lupus, B. taurus, M. musculus, R. norvegicus, G. gallus, D. rerio*.

#### 2.3.2 Prediction of RNA Secondary Structures

The expression of gene depends on the transcript stability and translational efficiency which can be affected by the modified secondary and tertiary structures of RNA. Using RNA structure web server (version 6.0.1), the secondary structure and stability of *BMP4* mRNA were predicted. This server employs algorithms based on thermodynamic principles to foretell the secondary structures of RNA. These predictions elucidate folding patterns. The sequence length of mRNA fragment comprises 41 nucleotides with the mutated nucleotide placed in the centre of the mRNA fragment. The change in minimum free energy was also noted using the same server for further statistical analysis. Further, the minimum free energy change (δδG) (δδG= δGMutant - δGWild type) was also calculated. The more positive is the δδG, the less stable is the mutated mRNA structure in comparison with the wild-type (WT) mRNA and vice-versa.

#### 2.3.3 Prediction of impact of mutants on RNA features

MutaRNA tool (version (5.0.10) was employed to further underscore the mutational analysis of the structural alterations induced by the mutated nucleotide [28]. The analysis involved the intra-molecular base pairing potential, base pairing probabilities of the mutant (MUT) mRNA, and RNA accessibility (single-strandedness) against the WT [29]. Changes induced by mutations in RNA structures can be statistically figured out by combining remuRNA and RNAsnp tools [30].

#### 2.3.4 Predicting the pathogenic potential of BMP4 novel variants

The disease-causing potential and the structural changes along with functional stability of proteins were predicted using Varcard2 (http://www.genemed.tech/varcards2/#/index/home) which is an integrated database of more than 15 predicting tools (SIFT, PROVEAN, PolyPhen-2, Mutation Taster, ClinPred, ReVe,, FATHMM, VEST3, LRT, CADD, GERP, DANN, and M-CAP etc.).

#### 2.3.5 Predicting the physiochemical properties of BMP4 proteins

The prediction to check the effect of variants on different physiochemical properties, a comparative analysis was performed for WT and BMP4 muteins. The changes in different physiochemical properties viz, relative mutability, ratio hetero end/side, alpha-helix and total beta strand were predicted using an *in silico* server, Protscale (https://web.expasy.org/protscale/).

#### 2.3.6 Structural alterations in the 2-D and 3-D conformation of BMP4 muteins

Psipred server (http://bioinf.cs.ucl.ac.uk/psipred/) was used to predict the structural changes caused by missense variants into the secondary structures of proteins. Moreover, the conformational changes into the tertiary structures were also predicted using the homology-based SWISS-MODEL server (https://swissmodel.expasy.org/interactive). The conformational alterations in the modelled-muteins over WT structures were visualized using UCSF Chimera which were further validated by global RMSD value, calculated by SuperPose 1.02 (http://superpose.wishartlab.com/).

### 2.4 In vitro analysis of BMP4 novel variants

#### 2.4.1 Cloning of BMP4 expression plasmid followed by site-directed mutagenesis

Human *BMP4* cDNA clone was purchased from “Addgene” (Cat No. #98321, Watertown, MA, USA). The protein coding region of BMP4, was PCR amplified to subcloned into pcDNAv3.1/NT-GFP TOPO TA vector (Invitrogen, Carlsbad, CA, USA), using PfuUltra high-fidelity DNA polymerase (Stratagene, Santa Clara, CA, USA), to prepare BMP4_WT_GFP. The mutant constructs (BMP4_R113G_GFP, BMP4_E151V_GFP, BMP4_T197I_GFP, BMP4_R226W_GFP) were prepared for all the identified variants (p.R113G, p.E151V, p.T197I, p.R226W) using Quick change II XL site-directed mutagenesis kit (Agilent Technologies Inc., USA) with a complementary pair of mega primers (as per manufacturer’s descriptions). The WT and mutant constructs were transformed into TOP10 competent cells followed by isolation of respective plasmids (Qiagen midi kit, Germany). Sanger sequencing was done to confirm the fidelity of all the constructs.

#### 2.4.2 Cell culture and transfection

To perform *in vitro* functional analysis, mouse pluripotent embryonic cell line, P19 (kind gift from Dr. Ramkumar Sambasivan, InStem, India) and cadiomyoblast H9c2 cell-line (purchased from cell repository, NCCS, India) were used. P19 cells were cultured in α-MEM (Gibco, Life technologies Corp.) while H9c2 cells were grown in DMEM, supplemented with 10% fetal bovine serum (Gibco, Life technologies Corp.) with 100 units/mL penicillin and 100 lg/mL streptomycin at 37°C with 5% CO2 in a humified chamber. For all *in vitro* experiments, ∼10^5^ cells were seeded in 6-well plates (for immunostaining, western blotting and qPCR) and 24-well plates (for luciferase assay). By using FuGENE 6 transfection reagent (Promega Corp., IN, USA) transfection was performed 24 hours after seeding, at 30-40% confluency.

#### 2.4.3 Estimation of expression and localization of BMP4 protein through immunostaining

Immunostaining was performed in H9c2 cell, due to their better morphology, in order to visualize the cellular localization of BMP4 WT and muteins. The cells were seeded in 6-well culture plate with each well containing glass cover slips and grown in DMEM, as described earlier. After 24 hours, BMP4_WT_GFP and MUT constructs (1µg) were transfected. Post 48 hours of transfection, cells were washed with chilled 1X PBS followed by fixation with 4% paraformaldehyde (PFA) for 15 minutes. For permeabilization, the fixed cells were treated with 0.5% Triton-X for 30 minutes. Cells were washed 3 times with chilled 1X PBS and stained with DAPI (Sigma-Aldrich, USA) and Phalloidin (Sigma-Aldrich, USA). Finally, mounted in mounting media and confocal scanning was performed using Leica SP8 STED Laser Scanning Super Resolution Microscope System, Germany. Images were analysed by ImageJ software and assembled by Adobe Photoshop software.

#### 2.4.4 Estimation of phospho-SMAD1/5 by Western Blotting

To investigate the expression of phospho-SMAD1/5 (Ser463/465) due to WT and MUTs, western blotting was performed. P19 cells were seeded followed by transfection with BMP4_WT_GFP and MUT constructs (1µg). Whole cell lysate was prepared in RIPA Buffer (as per Dixit et al, 2021) post 48 hours of transfection followed by quantification of proteins by Bradford assay. 25µg of reducing samples were denatured for 5 min at 95°C in Laemmeli’s loading buffer and lysates were resolved on 8% SDS polyacrylamide gel followed by transferring the proteins to PVDF membrane (Bio-Rad Laboratories Inc, CA, USA). After blocking the membrane in 5% non-fat dry milk in 1X TBST for 2 hours at RT, it was incubated with phospho-SMAD1/5 primary antibody (CST, USA) at 4°C for overnight followed by washing with 1X TBST (3 times) and then incubated with HRP conjugated goat anti-Rabbit IgG antibody (Genei, Germany) for 2 hours at RT. The membrane was washed with 1X TBST (5 times) and chemiluminescence detection was carried out with ECL detection kit (GE health Care, US) using chemiDoc (Amersham^TM^ Imager 680, Japan).

#### 2.4.5 Evaluation of transcriptional activity by Dual Luciferase Reporter Assay

The transactivation of BMP responsive promoters namely *Id1-luc* and *Id3-luc* due to BMP4 muteins was checked by dual luciferase reporter assay in P19 cells. Luciferase reporter plasmids i.e., *Id1-luc* and *Id3-luc,* used in the reporter assay were obtained as a kind gift from (Maria Genander, Karolinska Institutet, Stockholm, Sweden); and (Daniel J. Bernard, Quebec, Canada) respectively. P19 cells were transfected with BMP4_WT_GFP) and mutant constructs (125 ng each), along with of each reporter plasmids (125ng) and an internal control Renilla luciferase vector (25ng), at 50% confluency after 24 hours. Following 24 hours of transfection, cells were washed with chilled 1X PBS followed by lysates preparation in 1X passive lysis buffer (PLB). The activity of luciferase was measured by Synergy/ HTX multi-mode reader (BioTek Instruments, Inc., USA) using Dual-Luciferase Reporter Assay System, (Promega Corp, WI, USA). For both the reporter plasmids, three independent experiments were performed in triplicate for each sample (WT and MUTs) and the data were pooled by plotting as mean fold change with standard error of mean. The significance of the data was tested by one-way ANOVA followed by Dunnett’s post-hoc test.

#### 2.4.6 Quantifying the effect of BMP4 variants on downstream target by qPCR assay

Real-time PCR was performed in P19 cells to estimate the effect of MUTs over BMP4_WT_GFP constructs on the endogenous expression level of downstream target genes namely *Id1, Id3, Smad1, Smad5* and *Irx4.* The cells were grown in 6-well culture plates followed by transfection after 24 hours with WT and MUT constructs (1µg). Ater 48 hours of transfection, total cellular RNA was extracted using TRI Reagent (Merck, Germany) followed by quantification using NanoDrop Micro UV/Vis spectrophotometer (ThermoFisher Scientific, USA). The integrity of RNA was checked on 1% agarose gel followed by DNase I (ThermoFisher Scientific, USA) treatment. 2 µg of DNase I-digested RNA was used to prepare cDNA library using Revertaid First Strand cDNA synthesis kit (ThermoFisher Scientific, USA) and Random hexamers. The q-PCR assay was performed using real-time PCR machine (QuantStudio 5, Applied Biosystems, CA, USA) and SYBR^Tm^ green (Sigma-Aldrich, USA) as per manufacturer’s instruction. Three independent experiments were performed in triplicate and the acquired data were plotted as fold change with standard error of mean.

## 3. Results

### 3.1 Novel BMP4 genetic variants with their genotype-phenotype association

Genetic screening of the coding exons and flanking exon-intron boundary regions of *BMP4* in 285 sporadic cases of CHD by Sanger’s method, identified a total of four novel missense variants (c.337C>G; p.R113G, c.452A>T; p.E151V, c.590C>T; p.T197I, c.676C>T; p.R226W) in four unrelated cases in heterozygous condition with minor allele frequency (MAF) <0.01. These novel variants were not reported in any clinical database, therefore submitted to ClinVar database. The submission numbers allotted for each variant are as follows: c.337C>G; p.R113G (SCV005044984), c.452A>T; p.E151V (SCV005044985), c.590C>T; p.T197I (SCV005044986), c.676C>T; p.R226W (SCV005044987). All the four variants were identified in pro-peptide domain of BMP4. None of these variants were detected in 400 ethnically aged-matched control (800 chromosomes). One of the missense variants, (**p.R113G**) was identified in a male child diagnosed with complex CHD i.e., Transposition of great arteries (TGA), double outlet right ventricle (DORV), atrial septal defects (ASD) and ventricular septal defects (VSD). The second pro-peptide domain variant (**p.E151V**) was also associated with complex phenotype of total anomalous pulmonary venous return (TAPVR), ASD and VSD in a female patient. The third missense variant (**p.T197I**) was also found in a female proband with a rare phenotype i.e., hypoplastic left heart syndrome (HLHS) and ASD. The last variant (**p.R226W**) was detected in a child with VSD. Due to unavailability of parental samples, their genotyping was not performed. Besides these, we also detected one SNP (p.V152A; rs17563) in 43 cases. The amino acid change with coding positions of nucleotide, the clinical CHD phenotype and MAF are presented in **Table1**. The electropherogram of sequence representing the above listed missense variants and its position on domain of protein are shown in **Figure 1(a) and (b)**.

**Figure 1.**
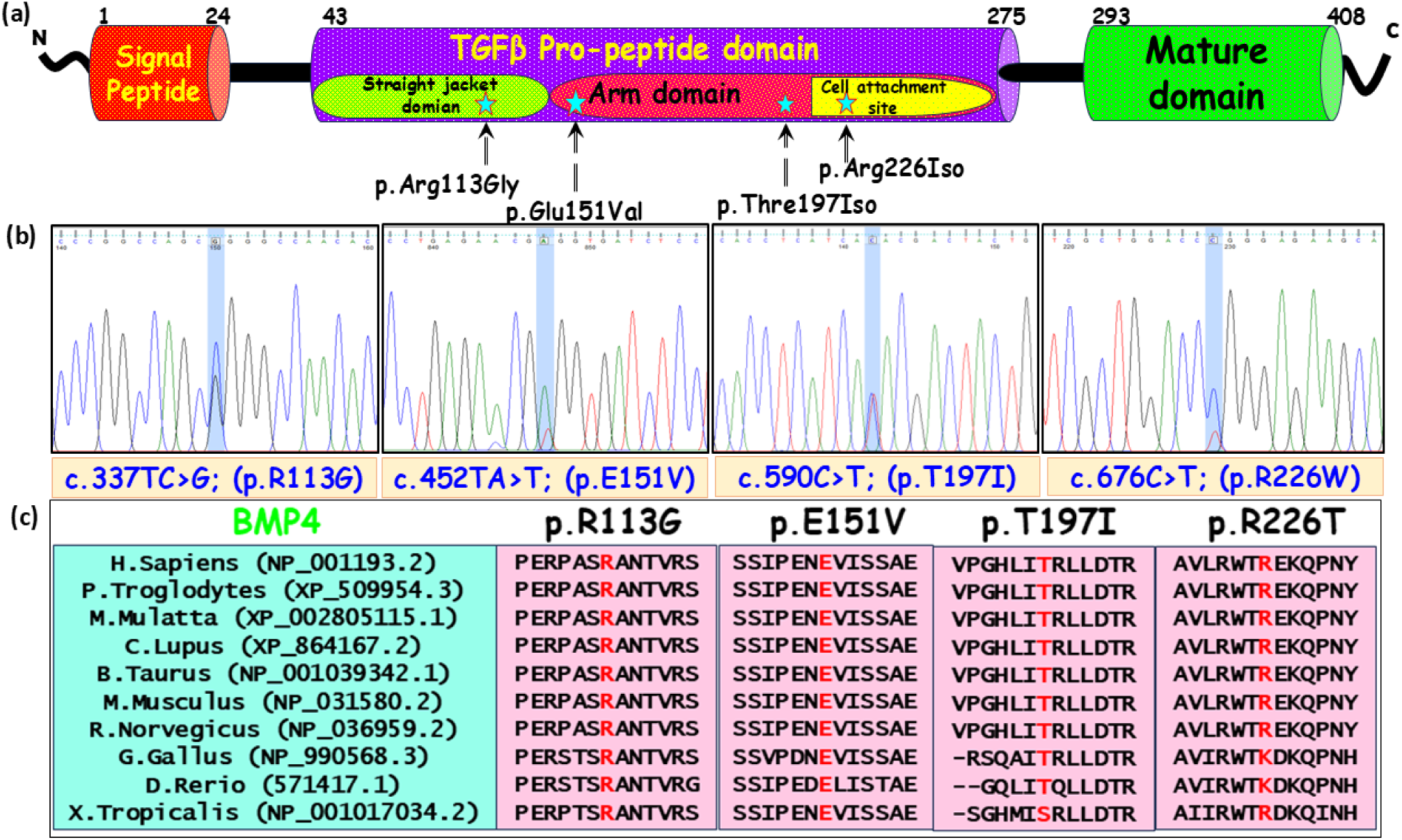
Diagrammatic illustration of BMP4 protein with identified variants and multiple sequence alignment. (a) Schematic representation of BMP4 protein with 408 amino acids representing different domains namely., signal peptide domain (1-24aa), TGFβ-Pro-peptide domain (43-275aa), protease cleavage site (276-292aa) and mature peptide domain (293-408aa). Domain-wise positioning of different identified missense variants (marked with star). (b) The sequence chromatogram for the identified variants and cross species alignment of the substituted amino acid across different vertebrate species (highlighted in red) for all the four variants (p.R113G, p.E151V, p.T197I and p.R226W).

**Table 1.** Clinical phenotypes, domain-wise variants position, ClinVar submission Ids and novelty status.

| Nucleotide change | AA change | CHD Phenotype | MAF | | BMP4 Domain | ClinVar | dbSNP | INDEX Database | Indigenome | 1000 Genome database | Genome Asia100K | gnomAD | I-mutant 2.0 ($\Delta\Delta G$, RI) |
| --- | --- | --- | --- | --- | --- | --- | --- | --- | --- | --- | --- | --- | --- |
|  |  |  | Cases (N=285) | Controls (N=400) |  |  |  |  |  |  |  |  |  |
| c.337C>G | p.Arg113Gly | dTGA, DORV, ASD, VSD | 1/285 (0.00175) | 0/285 | Pro-peptide | SCV005044984 | NR | NR | NR | NR | NR | NR | Decrease (-2.88, 8) |
| c.452A>T | p.Glu151Val | TAPVR, ASD, VSD | 2/285 (0.00351) | 0/285 | Pro-peptide | SCV005044985 | NR | NR | NR | NR | NR | NR | Increase (-0.72, 4) |
| c.590C>T | p.Thr197Ile | HLHS, ASD | 1/285 (0.00175) | 0/285 | Pro-peptide | SCV005044986 | NR | NR | NR | NR | NR | NR | Decrease (-0.25, 3) |
| c.676C>T | p.Arg226Tryp | VSD | 1/285 (0.00175) | 0/285 | Pro-peptide | SCV005044987 | NR | NR | NR | NR | NR | NR | Decrease (-1.74, 8) |

**Abbreviations:** dTGA- dextro Transposition of great arteries, DORV- double outlet right ventricles, ASD- Atrial septal defects, VSD- Ventricular septal defects, TAPVR- Total anomalous pulmonary venous return, HLHS- Hypoplastic left heart syndrome, MAF- Minor allele frequency, dbSNP- single nucleotide polymorphism database, NR- Not reported, RI- Reliability index

### 3.2 Evolutionary conserved BMP4 protein across different species

The phylogenetic conservation of the substituted amino acids was checked across different species of vertebrates namely, *Homo sapiens* (NP_001193.2), *Pan troglodytes* (XP_509954.3) *Macca mulatta* (XP_002805115.1) *Canis lupus* (XP_864167.2) *Bos taurus* (NP_001039342.1) *Mus musculus* (NP_031580.2) *Rattus norvegicus* (NP_036959.2) *Gallus gallus* (NP_990568.3) *Danio rerio* (571417.1) *Xenopus tropicalis* (NP_001017034.2) were checked. The amino acid residues are highly conserved at positions Arg113, Glu151, Thre197 and Arg226 across above listed vertebrate species. The conservation of the substituted amino acids residues along with the flanking amino acid sequences has been shown in **Figure 1(c)**.

### 3.3 Predicted pathogenic potential of BMP4 identified variants

The potential to cause disease by the variants were predicted by the VarCard2 server. Most of the tools SIFT, Polyphen2_HDIV, M_CAP, Mutation Taster, Eigen, GenoCanyon, CADD, and DANN predicted that all the four novel missense variants are damaging and disease causing. Table 2 represents the predictions and the algorithmic scores of all the four variants as predicted by different *in silico* tools **(Table 2).**

**Table 2.** Predicted disease-causing potential of BMP4 variants by different in-silico tools

| Variant | SIFT |  | Polyphen2_HDIV_pred |  | M_CAP_pred |  | Mutation Taster |  | Eigen |  | GenoCanyon_pred |  | CADD |  | DANN |  |
| --- | --- | --- | --- | --- | --- | --- | --- | --- | --- | --- | --- | --- | --- | --- | --- | --- |
|  | Score | Prediction | Score | Prediction | Score | Prediction | Score | Prediction | Score | Prediction | Score | Prediction | Score | Prediction | Score | Prediction |
| c.337C>G | 0.02 | Damaging | 1 | Probably_damaging | 0.068 | Damaging | 1 | Disease_causing | 1 | Damaging | 0.61 | Damaging | 21.8 | Damaging | 0.994 | Damaging |
| c.452A>T | 0 | Damaging | 1 | Probably_damaging | 0.193 | Damaging | 1 | Disease_causing | 1 | Damaging | 0.677 | Damaging | 27.2 | Damaging | 0.992 | Damaging |
| c.590C>T | 0.022 | Damaging | 0.853 | Possibly_damaging | 0.074 | Damaging | 1 | Disease_causing | 1 | Damaging | 0.677 | Damaging | 22.4 | Damaging | 0.99 | Damaging |
| c.676C>T | 0.016 | Damaging | 0.954 | Possibly_damaging | 0.054 | Damaging | 0.999 | Disease_causing | 1 | Damaging | 0.677 | Damaging | 29.9 | Damaging | 0.999 | Damaging |

**Abbreviations:** SIFT- Sorting Intolerant from tolerant, CADD- Combined annotation dependent depletion, DANN- Deleterious annotation using neural networks

### 3.4 Impact of BMP4 variants on RNA structures and stability

The secondary structural changes of mRNA were checked and the analysis results depicted prominent alterations in the structures due to three pro-peptide domain c.337C>G (p.R113G), c.452A>T (p.E151V), c.676C>T (p.R226W) variants against WT. The significant changes speculated to affect the RNA folding and stability. This observation was further supported by the change in free energy (δδG) which is 3.0kJ/mol, 2.0kJ/mol and −3.4kJ/mol due to variants c.337C>G (p.R113G), c.452A>T (p.E151V), c.676C>T (p.R226W) respectively. However, one of the pro-peptide domain c.590C>T (p.T197I) variant, exhibited less pronounced modifications in mutant structure compared to WT with change in free energy (δδG) was 0.0kJ/mol **(Figure 2a1-a4).**

**Figure 2.**
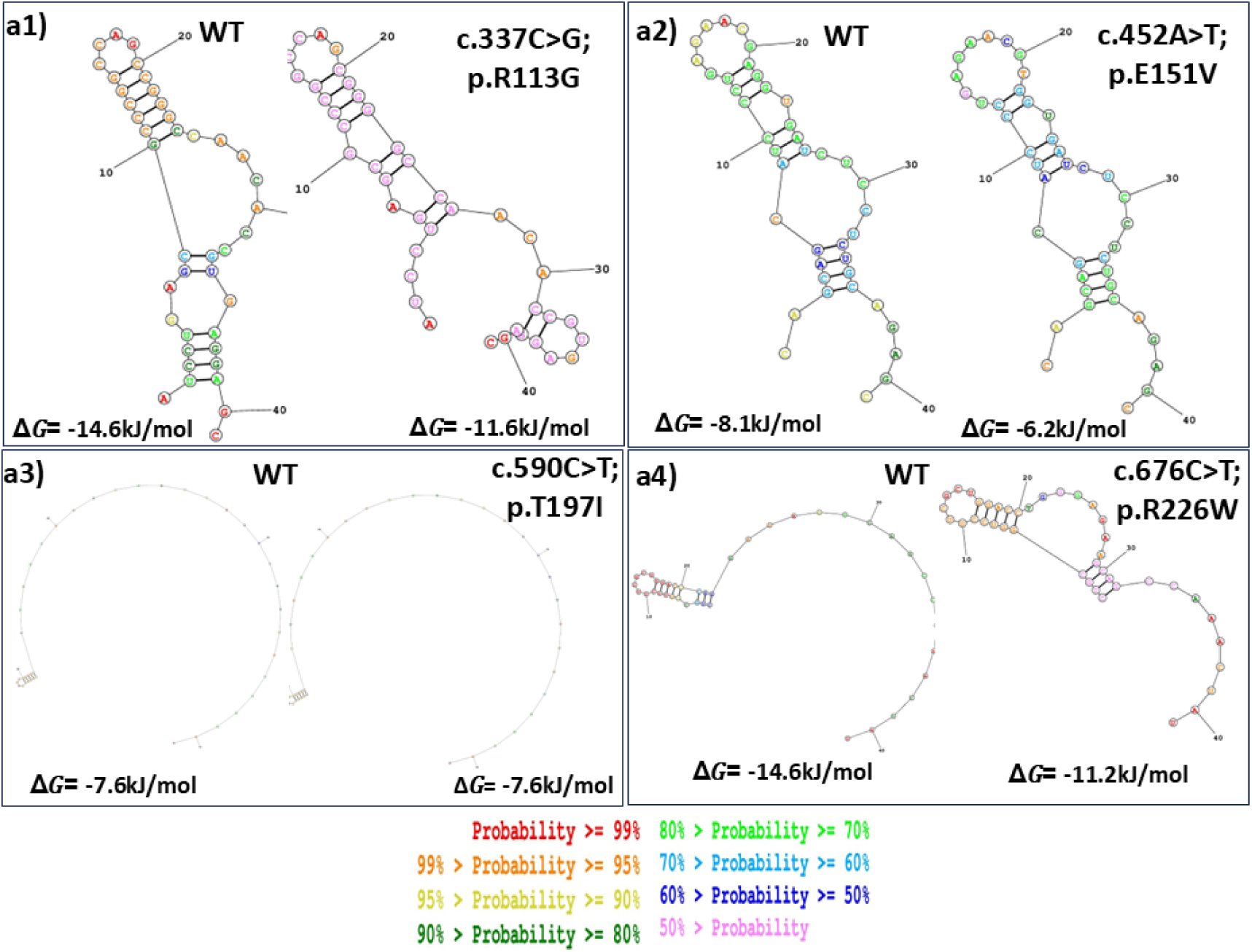
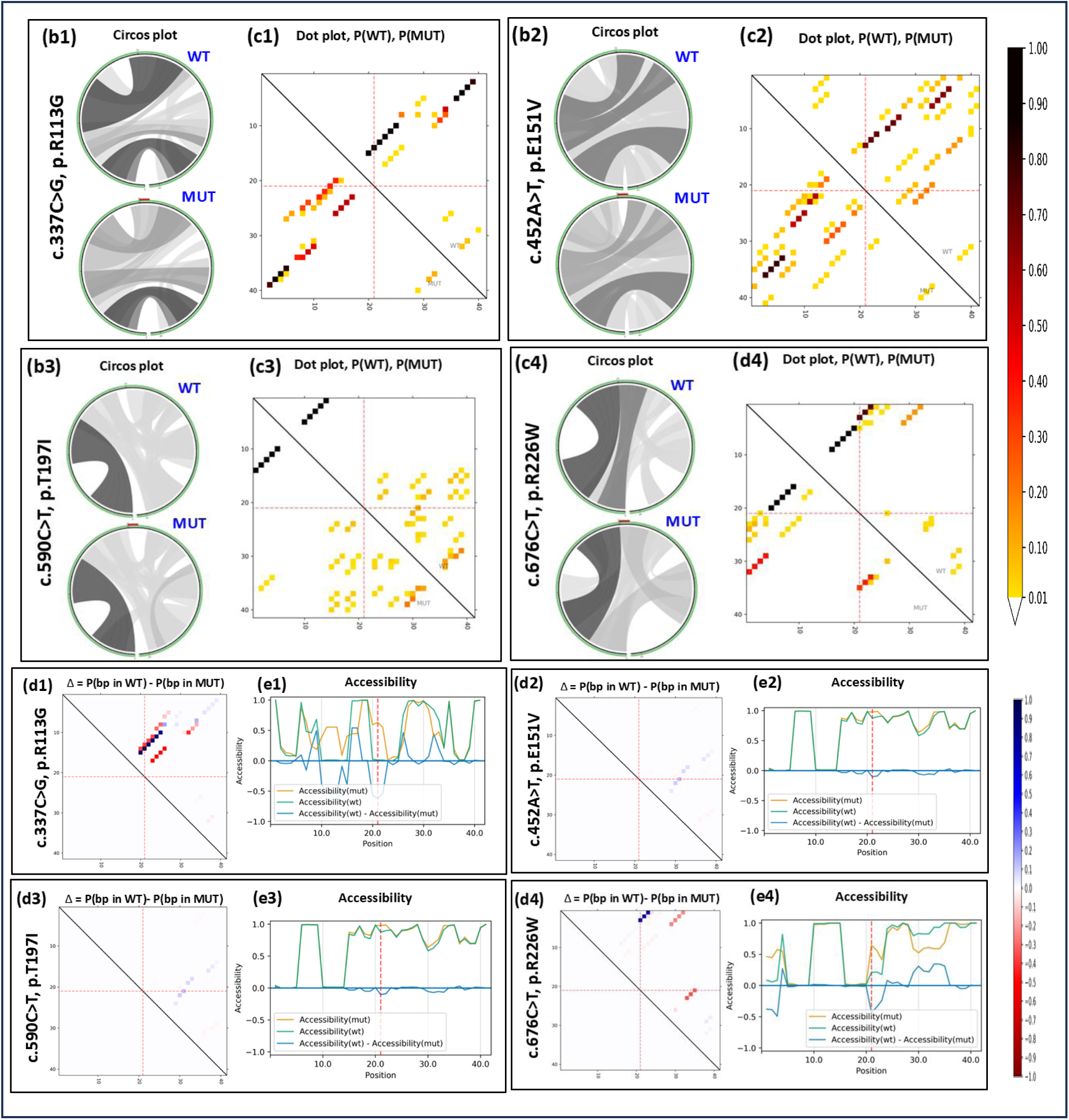
(a1-a4) Predicted secondary structures of WT and MUT RNA using RNA structure web server for all the four variants c.337C>G (p.R113G), c.452A>T (p.E151V), c.590C>T (p.T197I), c.676C>T (p.R226W) of BMP4. The WT structure is presented on left and MUT on right. The nucleotide change is at 21st position in each structure. Different colors indicating different base pairing probabilities. (b1-b4) Analysis of base pair probability by Circos and dot plot. The Circos plot illustrating the probability of base -pair for p(WT) and p(MUT) BMP4 RNA sequence, with the sequence initiating from 5’ end at the bottom-left and elongating clockwise reaching to the 3’ end. Each MUT Circos plot is highlighting the variant of interest in red at position 21 at the top. The darker shades of grey signify the base pairing of higher probability. (c1-c4) The heatmap-like dot plot representing base -pair probability of p(WT) and p(MUT) RNA variants. The darker dots symbolize more chance of the base WT sequence while the bottom left with the MUT sequence. The variant of interest is at position 21 highlighted by red dotted lines on both the axes. (d1-d4) The dot plot depicts the differences in base pairing probabilities between the WT and MUT RNA which is calculated as {Pr(bp in WT) – Pr(bp in mut)}. Weak base-pairing is represented by blue color dots and the red color dots are the results of strong base-pairing. (e1-e4) The accessibility profile of WT and MUT are denoted in terms of unpaired probabilities. The variation in accessibility (WT-MUT) is illustrated by the blue line.

### 3.5 Effect of variants on RNA structural features

Assessment on different features of RNA was portrayed by circos plot base pairing probabilities, base pairing probabilities dot plot, differential base pairing probabilities dot plot and accessibility profile to highlight the effect of variations. Circos plot represented the interconnection between different positions of RNA sequence and their interactions. The arcs which represent the variants connecting the affected nucleotides and illustrate the potential disruption caused by variants using different hues of grey, with darker grey color shows potentially stronger base pairing probabilities **(Figure 2b1-b4).** Similarly, the heat map dot plot matrices also indicate the base pairing probabilities with darker dot depicting higher base pairing potential **(Figure 2c1-c4)**. Further, the difference in base pair probability (Pr (bp in WT) – Pr (bp in MUT)) were also denoted by another dot plot matrices (differential dot plot) which define the difference in base pairing patterns between WT and MUT RNAs at specific locations. Visualised changes in the differential dot plot were observed in case of all the four variants with red dots demonstrating strong base pairs and blue dots less interaction. Remarkable changes were noted in all the four c.337C>G (p.R113G), c.452A>T (p.E151V), c.590C>T (p.T197I), c.676C>T (p.R226W) variants underscoring significant deviation in mutant RNA structure compared to WT counterpart **(Figure 2d1-d4)**. Additionally, the RNA accessibility profile which define the probability of being unpaired at each base position that affect the RNA-protein or RNA-RNA interactions, the crucial factors of translation was also explored. The analysis results indicating a significant difference in RNA accessibility due to all the c.337C>G (p.R113G), c.452A>T (p.E151V), c.590C>T (p.T197I), c.676C>T (p.R226W) variants **(Figure 2e1-e4).**

### 3.6 Deformities in 2-D and 3-D structures of BMP4 muteins

The altered secondary structures of mutant proteins (muteins) depicted significant changes in α-helix and β-sheet. There were shortening of α-helix at amino acid (aa) position 3-25 and 317-319, extension from 284-290aa position along with addition from 151-152aa position in mutein p.R113G. Additionally, shortening (201-205aa) and extension of β-sheet (315-319aa) was also noted in this variant. Similarly, shortening of α-helix at aa position 4-19 and extension at aa position 283-288 along with extension in β-sheet at aa 233-240 position for mutein p.E151V was portrayed. Moreover, α-helix was lost at three different positions from 29-30aa, 36-37aa and 123-125aa, extension in β-sheet from 233-240aa and 313-316aa was illustrated in mutein p.T197I. Similarly, shifting in α-helix at aa position 35-38, lost at 123-125aa position and extension at 266-267aa and 283-289aa position was demonstrated in p.R226W. Marked changes in β-sheet were also observed in this variant. All these alterations in mutein secondary structures were compared to WT **(Supplementary** Figure 1).

Furthermore, changes in the tertiary structure of muteins were visualized by superimposing mutein and WT structures by using UCSC Chimera. Considerable changes in the tertiary structures of muteins p.R113G and p.R226W was portrayed by the superimposed structures with supported RMSD value of 0.17 Å for both the muteins. However, no visualized changes in the tertiary structures of p.E151V and p.T197I were depicted **(Figure 3a-e).**

**Figure 3.**
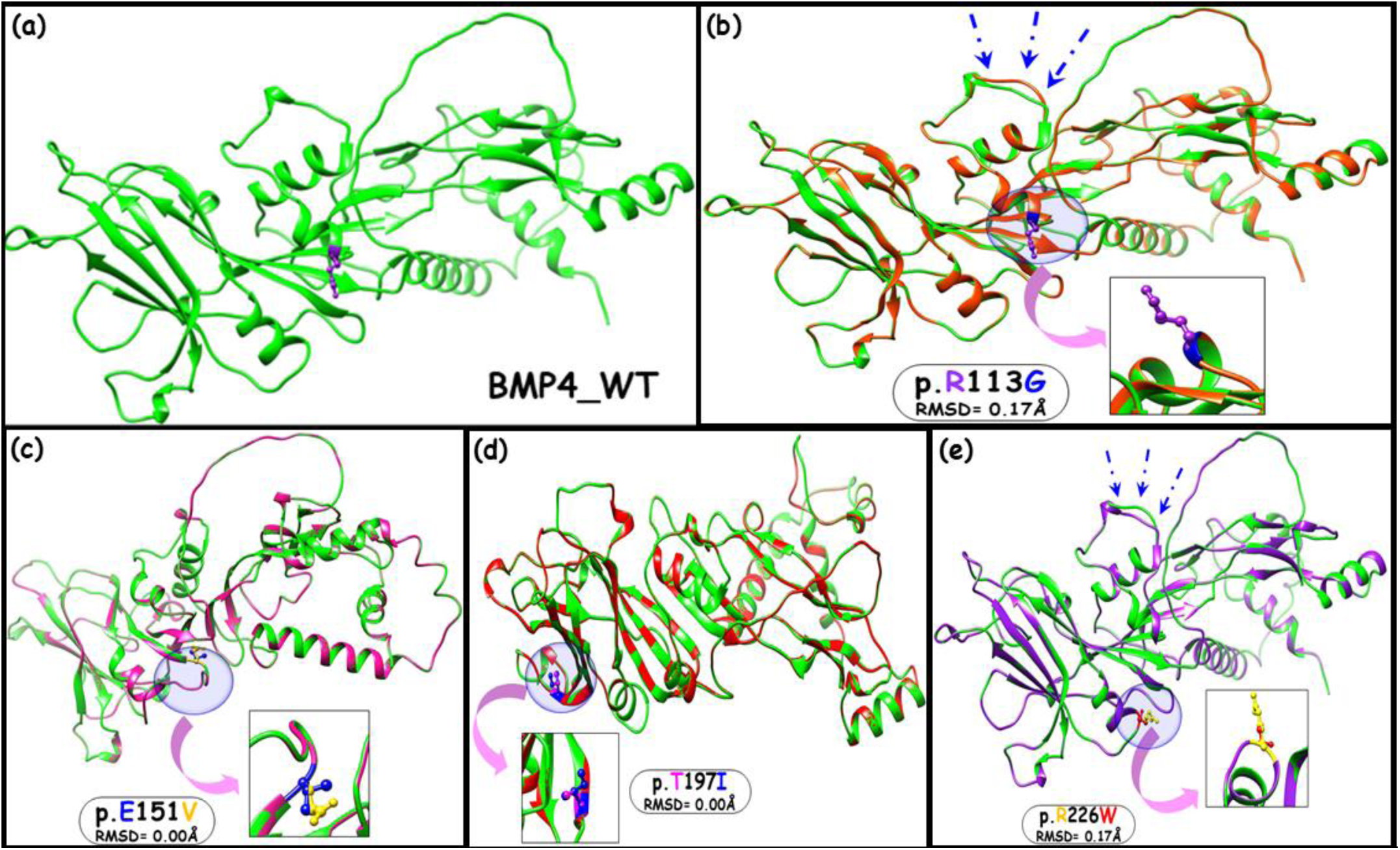
(a) Predicted tertiary structure of BMP4 WT (green colour), (b-e) Aligned tertiary structure of BMP4 WT against different muteins (p.R113G, p.E151V, p.T197I and p.R226W) in different colour. WT and mutated residues are illustrated by ball and stick model in different shades and regions having confirmational alterations in superimposed structures are marked by arrows.

### 3.7 Impact of variations in changing the physiochemical properties of BMP4 protein

The physiochemical properties of proteins depend on the R-side chain of amino-acids. The missense variants introduce new amino acid with different R-side chain which likely change the secondary and tertiary structures and functions of proteins. Different physiochemical properties such as relative mutability, ratio hetero end side, alpha-helix, and total -beta strand was predicted by Protscale for WT and muteins. Comparable changes were observed in relative mutability due to all the four (p.R113G, p.E151V, p.T197I, and p.R226W) variants. Similarly, the predicted analysis for ratio hetero-end were also depicted remarkable changes. Further, the alpha-helix and total beta-strands were also changed notably in all the four muteins (p.R113G, p.E151V, p.T197I, and p.R226W) **(Supplementary** Figure 2).

### 3.8 Effect of identified variations on the expression and localization of BMP4 protein

Immunofluorescence staining was conducted in H9c2 cells which demonstrated that BMP4_WT_GFP and all the four (p.R113G, p.E151V, p.T197I, and p.R226W) muteins tagged with GFP were localized to both cytoplasm as well as in nucleus. In all the muteins, protein aggregation in the form of larger spots were observed which were less defined in WT. However, there was no significant changes observed in the expression and localization of muteins when compared to the WT **(Figure 4a1-e4).**

**Figure 4.**
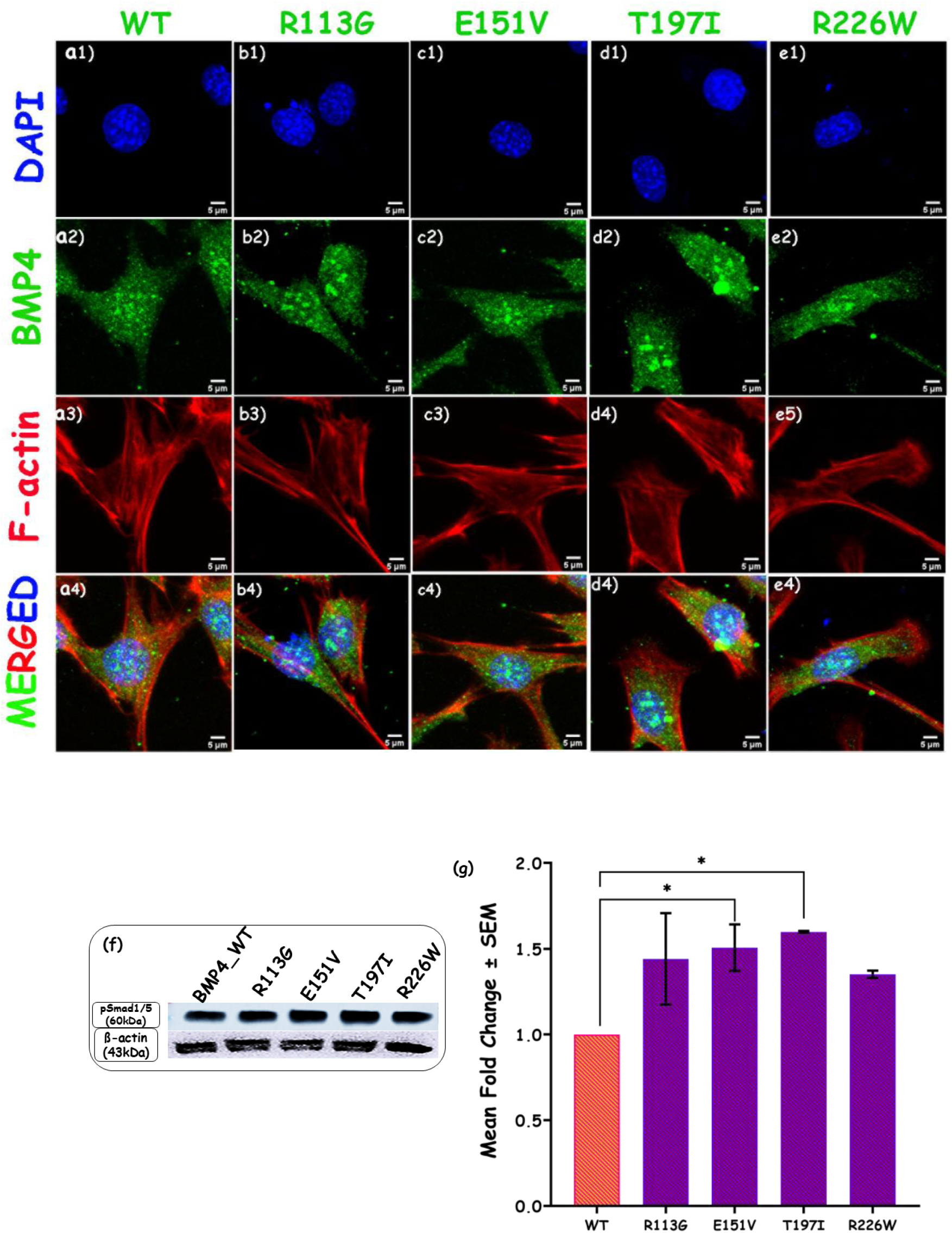
Expression analysis of BMP4 wild-type (WT) and muteins (MUT) by immunostaining and western blotting. (a1-e4) Immunostaining demonstrating cellular location of BMP4 WT and MUT (p.R113G, p.E151V, p.T197I and p.R226W) in H9c2 cells. (f & g) Western blotting showing phosphorylation of SMAD1/5 due to WT and all four variants (p.R113G, p.E151V, p.T197I and p.R226W).

### 3.9 Impact of missense variants on the expression of BMP4 and phospho-SMAD1/5 protein

The effect of BMP4 variants on the phosphorylation of SMAD1/5 was estimated in P19 cells by western blotting. The phosphorylation of SMAD1/5 was significantly enhanced by 1.507 (p = 0. 0.0370) and 1.599 (p = 0.0193) fold due to p.E151V and p.T197I variants respectively when compared to the WT **(Figure 4f-g)**. Nonetheless, the other two variants p.R113G, and p.R226W also increase the phosphorylation of SMAD1/5 by 1.441 (p = 0.0614), 1.351 (p = 0.1275) fold respectively though statistically not significant **(Figure 4f-g).**

### 3.10 BMP4 variants affect the basal transcriptional activity of Id1and Id3 promoters

The transactivation of downstream promoters of BMPs *viz, Id1-luc* and *Id3-luc* were checked by dual reporter luciferase assay in response to BMP4 muteins. The transcriptional activity of each promoter i.e., *Id1-luc* and *Id3-luc* were normalized with the basal activity of these promoters. The transcriptional activity of *Id1-luc* and *Id3-luc* was 2.83 fold (p=0.0676) and 6.34 fold (p = 0.0044) respectively in proportionate to BMP4 WT **(Figure 5a-b)**. Fold change was calculated as the activity of these promoters caused due to BMP4 MUTs while comparing to WT. The transcriptional activity of promoter *Id1-luc* was significantly increased due to three pro-peptide domain variants p.R113G (1.83 fold, p = 0.0167); p.T197I (1.89 fold, p = 0.0114) and p.R226W (1.76 fold, p = 0.0287) while, mutein p.E151V (1.57 fold, p = 0.1149) also upregulate the expression of *Id1-luc* though statistically not significant **(Figure 5a).** Further, transactivation of *Id3-luc* promoter also showed enhanced expression. Although in case of proximal variant, p.R113G (1.12 fold (p = 0.8789) the increase was marginal and not significant the other three distal variants p.E151V (1.72 fold; p=0.0093), p.T197I (1.54 fold; p = 0.0358), and p.R226W (1.55 fold; p = 0.0332) showed significant upregulation **(Figure 5b).**

**Figure 5.**
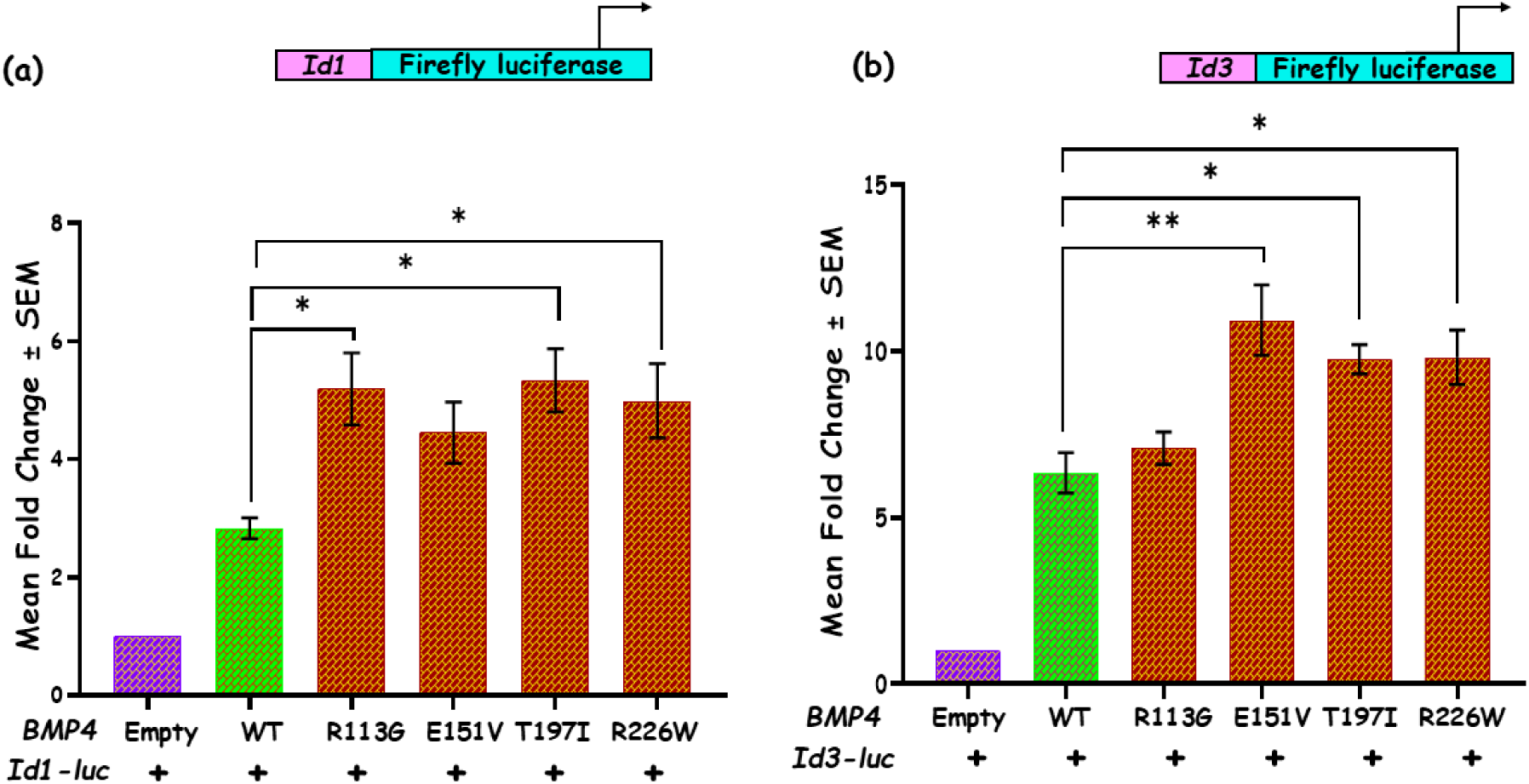
Transcriptional activity of *Id1-luc* and *Id3-luc* measured by dual luciferase reporter assay (a) Transactivation of *Id1-luc* due to BMP4 WT and MUTs (p.R113G, p.E151V, p.T197I and p.R226W) (b) Transcriptional activity of *Id3-luc* in response to BMP4 WT and MUTs (p.R113G, p.E151V, p.T197I and p.R226W).

### 3.11 Effect of missense variants on the expression of downstream target genes of BMP4

The effect of MUTs on the endogenous expression of transcription factors (*Id1*, *Id3* and *Irx4*) and co-transcriptional activators (*Smad1* and *Smad5*) of BMP signaling was also investigated by real-time PCR. The expression of one of direct target of Smads, namely *Id1* was significantly increased in response to all the four-novel p.R113G (1.372 fold; p < 0.0001), p.E151V (1.314 fold; p = 0.0003), p.T197I (1.255 fold; p = 0.0033), and p.R226W (1.55 fold; p < 0.0001) variants **(Figure 6a)**. Likewise, notable enhancement in the expression of another transcription factor *Id3,* was also depicted. Significant up-regulation was observed by three p.R113G (1.316 fold; p < 0.0001), p.E151V (1.385 fold; p < 0.0001), and p.R226W (1.276 fold; p < 0.0001) variants while, variant p.T197I (1.137 fold; p = 0.0861) although showed similar trend in increase in the expression of *Id3,* however not significantly **(Figure 6b).** Further, a significant upregulation of *Irx4* was also observed due to all the four p.R113G (1.525 fold; p < 0.0001), p.E151V (1.322 fold; p = 0.0023), p.T197I (1.289 fold; p = 0.0067), and p.R226W (1.362 fold; p = 0.0006) variants **(Figure 6c).** Additionally, when the expression of co-transcriptional activators was checked, a similar significant increase in the expression of *Smad1* was observed in response to all the p.R113G (1.227 fold; p < 0.0001), p.E151V (1.238 fold; p < 0.0001), p.T197I (1.144 fold; p = 0.0107), and p.R226W (1.177 fold; p = 0.0015) variants **(Figure 6d).** Moreover, a significant increment in the expression of *Smad5* was also exhibited due to p.E151V (1.196 fold; p = 0.0002), p.T197I (1.546 fold; p < 0.0001), and p.R226W (1.302 fold; p < 0.0001) variants, except for p.R113G variant, in which enhanced the expression of Smad5 was 1.102 fold (p = 0.0786), albeit not statistically significant **(Figure 6e)**.

**Figure 6.**
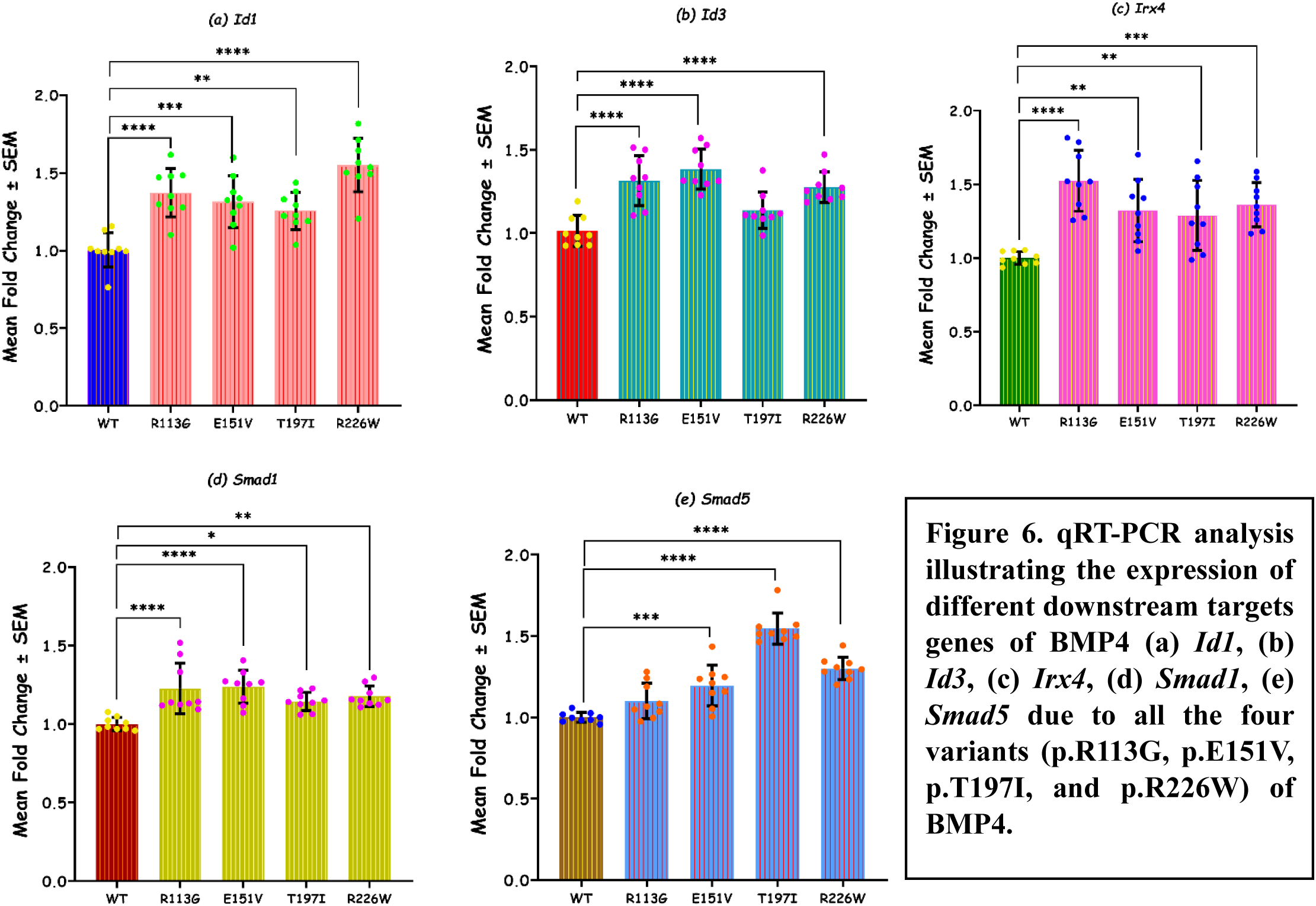
qRT-PCR analysis illustrating the expression of different downstream targets genes of BMP4 (a) *Id1*, (b) *Id3*, (c) *Irx4*, (d) *Smad1*, (e) *Smad5* due to all the four variants (p.R113G, p.E151V, p.T197I, and p.R226W) of BMP4

## 4. Discussion

BMP4 is one of the critical signaling molecule plays important role during embryonic patterning and axis formation, predominantly express in ventral side of the embryo, controlling dorsal and lateral mesoderm and neural plate formation [31,32]. BMP4 is a 408 aa long protein (44.04KDa), that comprises a signal peptide (24 aa), a pro-peptide domain (232 aa), and a mature peptide domain (115aa). Genomic location of human *BMP4* is on chromosome 14q22.2 (HGNC:1071), spanning approximately 7 kb, encoded by 6 exons, however exon 3 and exon 4 are the only protein coding exons. The protein form homo/heterodimers to trigger type II receptor (BMPRII) that activate type I receptor (BMPR1A) resulting in the phosphorylation of downstream SMADs (SMAD1/5 and SMAD4) transducers [18,31]. The SMAD complexes bind and regulate a diverse group of cardiac enrich transcription factors viz., *Id1* [32–34], *Id2* [35], *Nkx2.5* [18,36,37] and *Gata4* [38,39]. Besides this, in a SMAD independent non-canonical signaling, BMP4 also activates MAP kinase pathway via TAK-1, a serine–threonine kinase [7,40]. Additionally, it is also involved in PI3K/Akt, Rho-GTPases pathways [7].

In the present study, we have identified four missense variants in 285 isolated probands of CHD which are absent in 400 healthy age-matched controls. All the four variants are novel as these are not reported in any of the genome variation databases viz. gnomAD, dbSNP, clinvar, 100G, GenomeAsia100K, INDEX-db, IndiGenomes databases. All the four variants lie in the pro-peptide domain of BMP4. The first (p.R113G) variant is reported in complex form of CHD phenotype (TGA, DORV, ASD, VSD), the second (p.E151V) is also associated with complex phenotype i.e., TAPVR, ASD and VSD. The third (p.T197I) is diagnosed in HLHS, ASD, while the fourth distal pro-peptide domain variants (p.R226W) is detected with VSD. Majority of phenotypes recognized in our study are vessels and septal defects. Such lesions are speculated to be possibly originated due to perturbation during endocardial and outflow tract cushion formation. In human, BMP4 has been previously reported as a candidate gene for CHD along with other developmental defects, however, majority of studies lacking the functional assessment of the identified variants. In 2007, Posch and his group [19] have shown the association of BMP4 SNP (rs17563, p.V152A) with congenital septal defects which is later demonstrated by Li et al, 2016 [21] to be significantly associated with CHD by statistical analysis. The same polymorphism (rs17563) has been also significantly associated with non-syndromic cleft lip with or without cleft palate (NSCLP) [41] and with subepithelial, microform, and overt cleft lip [24]. In the current investigation we also detected SNP (rs17563, p.V152A) in 43 CHD cases with diverse phenotype. Recently in a Chinese study, a non-sense variant (Y106X) is identified in pro-peptide domain of BMP4 which reduced the transcriptional activity of *NKX2-5* and *TBX20* promoters [22]. Additionally, loss-of-function mutations in BMP4 are also reported in developmental eye disorders including SHORT syndrome [25] and in tooth agenesis and low bone mass [27] as well as combined pituitary hormone deficiency [42]. Moreover, previous reports on abnormal cardiac development and embryonic lethality in Bmp4 knockout mouse [43] as well as septal defect particularly VSD in murine embryos double heterozygous for Bmp4 and Bmp2 [9,44] has suggested critical role of BMPs during endocardial cushion formation and septation. Moreover, conditional deletion of Bmp4 in cardiomyocyte in mice also results in VSD, atrioventricular canal defect (AVCD) and DORV [17]. Likewise, conditional deletion of Bmpr2 (BMP receptor) gene in mice leads to VSD, DORV and AV cushion defects which further validating the role of BMP signaling in chamber myocardium [48]. Interestingly, substantial investigations, reporting many downstream genes of BMP signaling as candidate gene for CHD namely *ALK2* [45], *ALK3* [46]*, BMPRII* [47,48], *SMAD1* [49], *SMAD4* [50], *GATA4* [51,52], *NKX2.5* [53], *IRX4* [54] are well-studied and known as crucial players for cardiogenesis. In a collaborative study, Wang et al., 2005 [55] have shown that *Alk2* mutant mice exhibited atrioventricular septal and valve defects [55]. A deleterious mutation (p.R443H) in *ALK3* has been identified in a familial CHD case [46]. Many pathogenic variants were identified in *BMPRII* which are linked with CHD [47]. Similarly, another pathogenic variant (p.A935V) was also identified in BMPRII in association with pulmonary atresia (PA) [48]. Wang and his group identified a non-sense variant in *SMAD1* in isolated CHD [49]. Likewise, SMAD4 also harbouring a non-sense variant investigated in isolated CHD [50]. Several studies reported disease-causing mutations in *GATA4* which are associated with isolated CHD [51,52]. Moreover, many reports available showing a large number of pathogenic variations in the master transcription factor *NKX2.5* [53,56]. In Chinese population, two pathogenic variants were reported in *IRX4* in non-syndromic CHD [54]. The null mice of Id1 exhibit valvular defects and outflow tract atresia as Id1 is a crucial gene for endocardial EMT in both chick and mouse embryo [57]. Id proteins are major players in early development and have crucial role in the differentiation and proliferation of cardiac progenitor cells and mature cardiomyocytes at multiple stages during heart development [31]. Additionally, the double or triple Id knockout embryos (*Id1*/*Id2*, *Id2*/*Id3*, *Id1/Id3* or *Id1*/*Id2*/*Id3*) demonstrate severe cardiac defects including valvular and septal defects, outflow tract atresia, impaired ventricular trabeculation and thinning of the compact myocardium layers; the embryos die at mid-gestation [58]. In BMPR1a-cKO hearts, the expression of Id1/3 was absent from the embryonic atrioventricular canal (AVC) which affects the formation of the atrioventricular valves and adjacent septa [59]. Furthermore, evidences from Noggin and Smad6 knockout mice, which lack endogenous BMP antagonism, exhibit boosted BMP signaling e.g., demonstrate valve hyperplasia, thereby reinforcing the significance of tightly regulated BMP4 activity in the pathogenesis of CHD [60]. The spectrum of cardiac phenotypes associated with BMP4 dysregulation likely reflects its fundamental involvement in key stages of heart development, including valvulogenesis, outflow tract morphogenesis, and endocardial cushion formation.

Functional validation of the rare variants observed in BMP4, using dual luciferase assay has depicted a significant increment in the transactivation of promoter *Id1-luc* caused by three p.R113G, p.T197I and p.R226W variants. The variant p.E151V has also shown 1.56 fold increase in the transcriptional activity of *Id1-luc* though could not meet statistical significance level. Similarly, *Id3-luc,* another target of Smad-complexes, also exhibit significant increase in the transcriptional activity due to three (p.E151V, p.T197I and p.R226W) variants, while p.R113G variant, show a marginal increase in the transactivation of *Id3-luc*.

Likewise, the impacts of all the four variants on the endogenous expression of different downstream targets of BMP4 namely *Id1*, *Id3*, *Smad1*, *Smad5* and *Irx4* has revealed overall enhancement in their expression levels. Among these, *Id1*, *Irx4*, and *Smad1* have exhibited a significant increase in the expression pattern due to all the four (p.R113G, p.E151V, p.T197I and p.R226W) variants. At the same time, although significant enhancement in the expression of *Id3* is noted due to three (p.R113G, p.E151V, and p.R226W) variants, the variant p.T197I has also showed notable increase, albeit not to a significant level. Similarly, the expression pattern of Smad5 has also significantly increased except for p.R113G variant. Taken together, our results from luciferase reporter assay as well as from real time downstream gene expression study demonstrate gain-of-function activity of these pro-domain variants. Smad-complexes directly regulate the expression level of *Id1* and *Id3* which further induce multiple downstream genes namely *Nkx2.5*, *Gata4* and *Irx4* in P19 cells [31] In a recent study, (2024) [22] have shown that a pro-domain nonsense mutation (Tyr 106X) causing depletion of *NKX2-5* and *TBX20* expression, however after transfection with a relatively high concentration of mutant constructs in HeLa cells which is not consistent with our observations[22]. Our observation in both real time qRT PCR and luciferase reporter assay clearly demonstrates gain-of-function activity.

Furthermore, we have also observed a remarkable increase in the phosphorylation of SMAD1/5 in response to all four variants, detected by Western blotting. Despite the fact that, two of these variants (p.E151V and p.T197I) show significantly high enhancement while other two variants (p.R113G and p.R226W) induce considerable higher fold increase in the phosphorylation status of SMAD1/5 albeit not statistically significant. It is well known that like other BMPs, BMP4 transduce signaling through SMAD dependent pathway, particularly *Smad1* and *Smad5,* which activate multiple downstream transcription factors [18,31]. In a recent study, ample expression level of BMP4 has been detected in normal and CHD-affected hearts without much difference [61]. However, they observed sustained mRNA and protein expressions in CHD patients [61]. Our previous study, using autopsy tissue samples from CHD heart, the expression pattern of BMP4 remain unchanged, when compared with control heart samples (data not shown, unpublished results) estimated by PCR-array (SAbiosciences, USA). The augmentated SMAD1/5 phosphorylation is likely dependent on the efficient binding of BMP4 mature ligand with its receptors, which further transduce the SMAD signaling. Further enhanced cleavage of mature-BMP4 is liable to pH and conformational changes in pro-domain and the protein folding that facilitate the cleavage [62,63]. Substitutions in p.R113G (arginine to glycine), p.E151V (glutamate to valine), p.T197I (threonine to isoleucine), and p.R226W (arginine to tryptophan) in the pro-domain are speculated to expose the PACE cleavage site, enhancing the intracellular processing of the BMP4 pro-domain. This enhanced processing likely leads to effective cleavage, secretion of mature BMP4, which in turn propitiously binds to the receptors and boosts SMAD-mediated signaling.

Immunostaining of BMP4 in H9c2 cells transfected with WT versus mutant constructs demonstrated both nuclear and cytoplasmic localization of the protein. However, careful observation indicates higher accumulation, blebbing of muteins compared to WT in the cells, which suggest possible delay in the turnover of mutant protein.

To further support our *in vitro* experiments, different *in silico* analyses are explored. Multiple sequence alignment highlights that the mutated amino acid residues are evolutionary conserved, which predict that the change in the amino acid may bring in changes in protein folding and stability, thus contributing to the activity of the protein which may contribute to pathogenic property of the variants [64]. Moreover, Varcard analysis predicted the disease-causing and damaging effect of all the four (p.R113G, p.E151V, p.T197I and p.R226W) variants. The pathogenic role of these variants underscores the association of these variants with CHD.

In addition to these, significant structural changes in mRNA structures due to variants p.R113G, and p.E151V has been illustrated. Nonetheless, remarkable alterations in other features of mRNA (the base pairing probabilities and accessibility profiles) induced by all the four variants (p.R113G, p.E151V, p.T197I and p.R226W), signify the damaging effect on RNA stability, function and RNA-protein interactions, the necessary factors required during developmental stages and helpful for diving into the molecular processes of disease pathogenesis [30].

Similarly, the physiochemical property analysis revealed remarkable changes in relative mutability, ratio hetero end side, alpha-helix, and total -beta strand of all the four muteins (p.R113G, p.E151V, p.T197I and p.R226W). These changes are likely to influence the protein’s overall structure, folding behaviour, stability, and interactions with other molecules [65]. In particular, modifications in secondary structure elements, including α-helices and β-sheets, indicate that these mutations may disrupt normal protein folding and function. Adjustments in α-helices could alter the protein’s shape and interfere with essential molecular interactions, while β-sheet disruptions may compromise the protein’s three-dimensional integrity and lead to aggregation. In addition, tertiary structure analysis revealed noticeable conformational shifts in muteins especially in p.R113G and p.R226W which showed distinct changes in helical and strand regions that could impair protein function. Thus, the computational and biochemical analyses suggest that these mutations alter the physiochemical properties and the secondary and tertiary structure of the protein, potentially increasing its interaction with binding partners through allosteric effects [66] further boosting BMP signaling.

## 5. Conclusion

Altogether, this study clearly demonstrates that gain-of-function mutations in pro-domain of BMP4 are associated with CHD phenotype. Our findings, supported by both *in vitro* and *in silico* data, point to upregulation of the BMP4 signaling that act as a contributing factor for the development of CHD. This supports the hypothesis proposed by Campos-Baptista (2008) that both loss- and gain-of-function mutations in BMP pathway components may underlie various cardiac anomalies [67]. Furthermore, gain-of-function mutations in other genes within the BMP signaling pathway have also been associated with CHD. Notably, variants in *TGFβ1* [68] and *CITED2* [69] have been linked to structural cardiac anomalies. Transcription factors such as *TBX20*, implicated in a ASD and valve malformations, have been shown to enhance the transcriptional activity of downstream targets [70]. Similarly, Postma et al. (2008) identified a novel *TBX5* variant, in a family with atypical Holt-Oram Syndrome that resulted in elevated TBX5-mediated transcription, further emphasizing the critical role of transcriptional regulation in cardiac development [71]. Notably, BMP signaling interacts with other key cardiac developmental pathways such as Nodal and TGF-β [13,60], NOTCH [13,36], and Wnt/β-catenin [72,73]. Upregulation of BMP signaling may interfere with these signaling networks, potentially leading to abnormal heart development. Further, evidence from Shi et al. (2016) [74] showed that alcohol exposure can enhance BMP target gene expression, suggesting environmental factors may also contribute to CHD risk through BMP pathway activation [74].

In a nutshell, our study highlights *BMP4* as a candidate gene for the development of human CHD with various pathogenic variants functionally associated with diverse cardiac phenotype. The functional characterization of variants uncovers previously unrecognized molecular mechanism involved in the pathogenesis of CHD. Varied phenotypes of CHD exhibited by *BMP4* variants would shed light in defining the role of *BMP4* at different stages of cardiogenesis.

## ACKNOWLEDGMENT

We are very thankful to all the patients, their family members and control individuals for their participation in the present study. We are extremely grateful to Prof. T.K. Lahiri and Damyanti Agrawal from Department of Cardio-vascular and Thoracic Surgery, IMS, BHU, Varanasi for their constant support and encouragement in enrolment of patients. We acknowledge University Grants Commission (UGC) for research fellowship (JRF and SRF) to Jyoti Maddhesiya.

## DATA AVAILABILITY STATEMENT

Raw data and derived data supporting the findings of this study are available from the corresponding author on request.

**Author Contribution**

Jyoti Maddhesiya performed all the *in silico* and *in vitro* experiments analyses and prepared the original manuscript draft, including all figures and tables. Ritu Dixit conducted the mutational screening. Prof. Bhagyalaxmi Mohapatra conceptualized the study, secured funding, and provided continuous oversight and revisions throughout the manuscript preparation. Prof. Ashok Kumar were responsible for the clinical evaluation and diagnosis of the patients enrolled in the study and assisted in collecting blood samples from patients and their available relatives. All authors reviewed and approved the final version of the manuscript.

## List of abbreviations

CHD: Congenital heart disease
ASD: Atrial septal defects
VSD: Ventricular septal defects
TGA: Transposition of great arteries
DORV: Double outlet right ventricle
TAPVR: Total anomalous pulmonary venous return
HLHS: Hypoplastic left heart syndrome
AVCD: atrioventricular canal defect
EMT: Epithelial to Mesenchymal Transformation

---

**Supplementary Table1.**
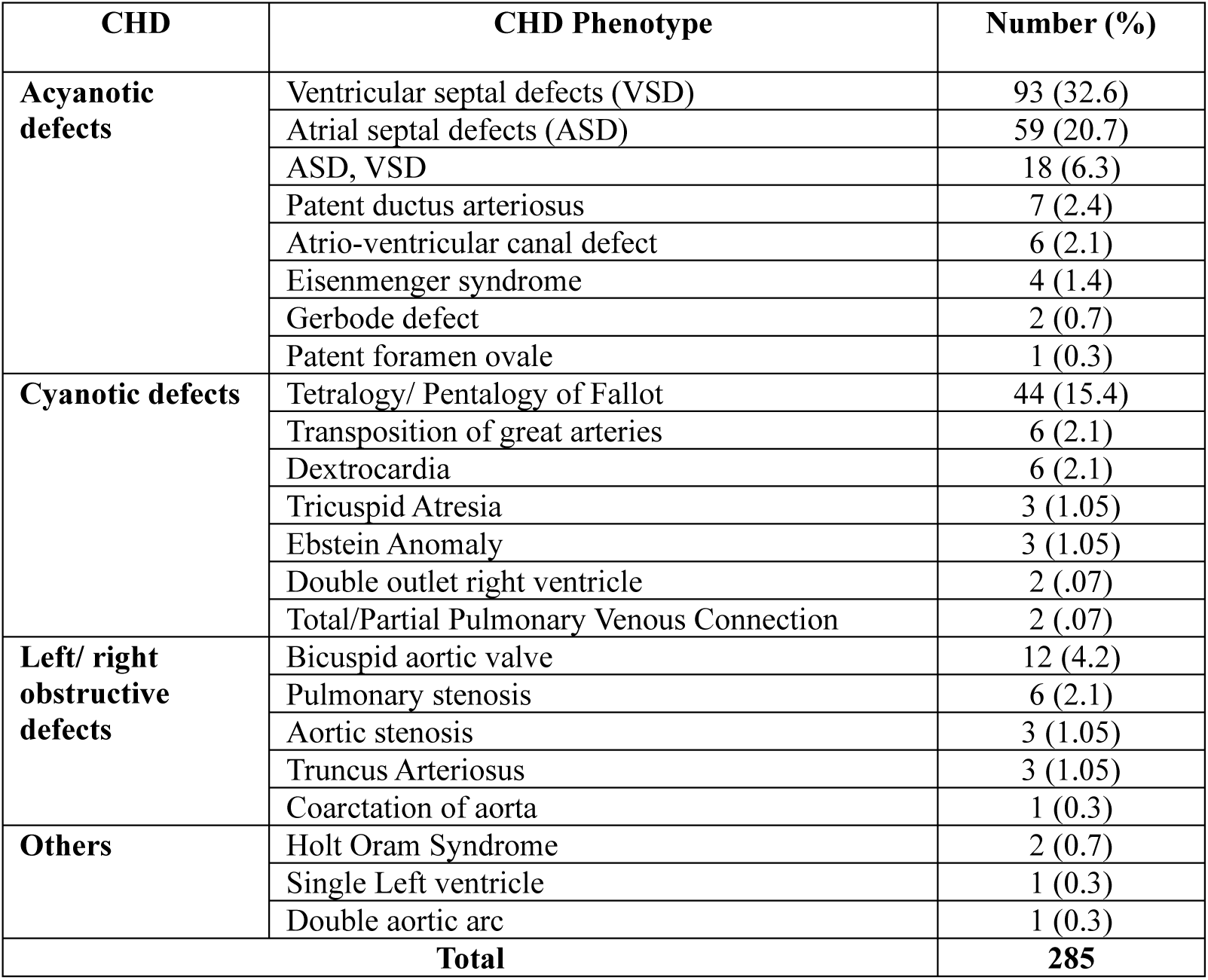
Clinical phenotype with their relative frequency in male and females.

| **CHD** | **CHD Phenotype** | **Number (%)** |
| --- | --- | --- |
| **Acyanotic defects** | Ventricular septal defects (VSD) | 93 (32.6) |
|  | Atrial septal defects (ASD) | 59 (20.7) |
|  | ASD, VSD | 18 (6.3) |
|  | Patent ductus arteriosus | 7 (2.4) |
|  | Atrio-ventricular canal defect | 6 (2.1) |
|  | Eisenmenger syndrome | 4 (1.4) |
|  | Gerbode defect | 2 (0.7) |
|  | Patent foramen ovale | 1 (0.3) |
| **Cyanotic defects** | Tetralogy/ Pentalogy of Fallot | 44 (15.4) |
|  | Transposition of great arteries | 6 (2.1) |
|  | Dextrocardia | 6 (2.1) |
|  | Tricuspid Atresia | 3 (1.05) |
|  | Ebstein Anomaly | 3 (1.05) |
|  | Double outlet right ventricle | 2 (.07) |
|  | Total/Partial Pulmonary Venous Connection | 2 (.07) |
| **Left/ right obstructive defects** | Bicuspid aortic valve | 12 (4.2) |
|  | Pulmonary stenosis | 6 (2.1) |
|  | Aortic stenosis | 3 (1.05) |
|  | Truncus Arteriosus | 3 (1.05) |
|  | Coarctation of aorta | 1 (0.3) |
| **Others** | Holt Oram Syndrome | 2 (0.7) |
|  | Single Left ventricle | 1 (0.3) |
|  | Double aortic arc | 1 (0.3) |
| **Total** |  | **285** |

