## Supplemental Figure for "Functionally significant, novel variants of *BMP4* are associated with isolated congenital heart disease"

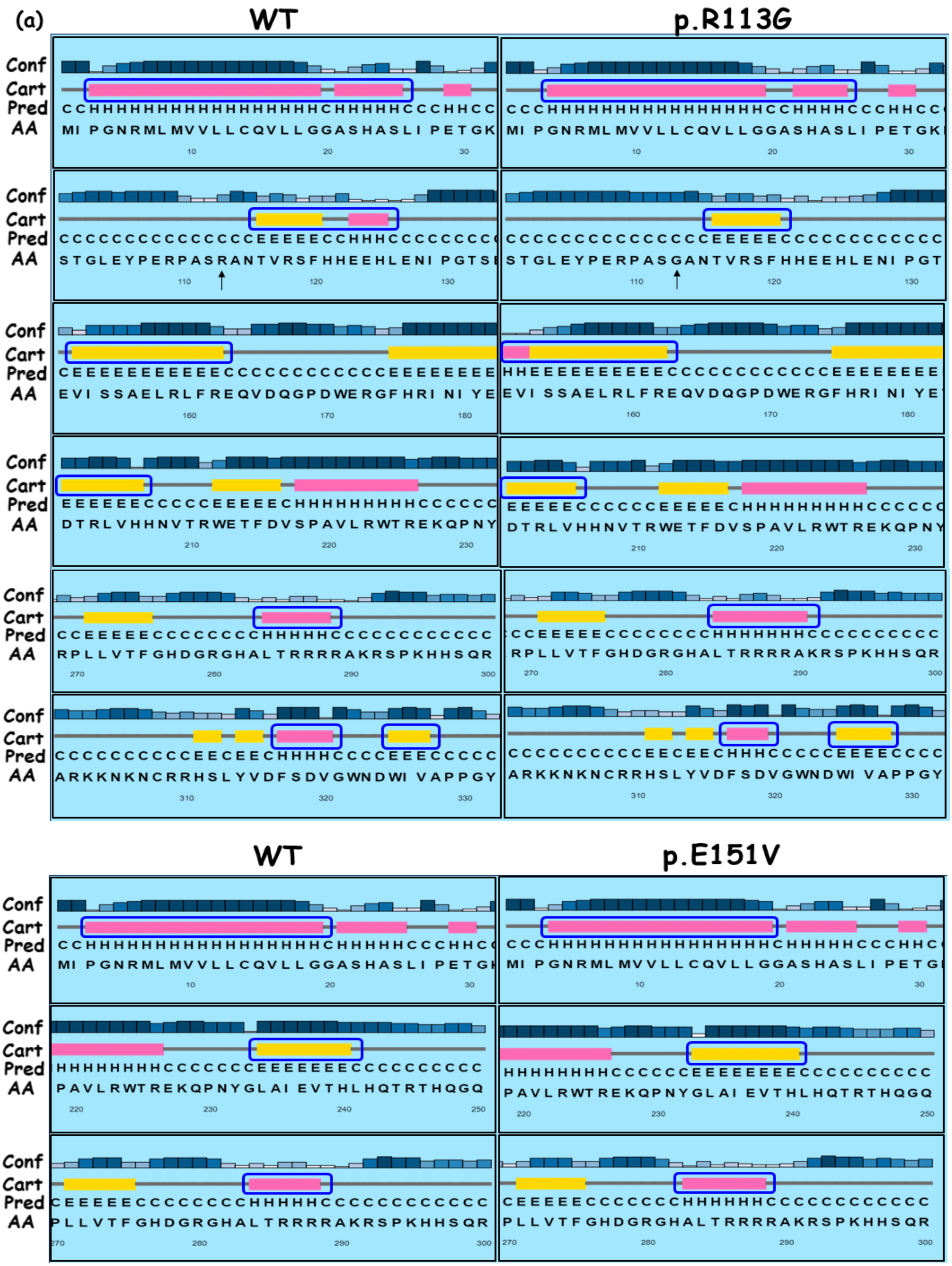
(a)
WT
p.R113G
Conf
Cart
Pred
AA
WT
p.E151V

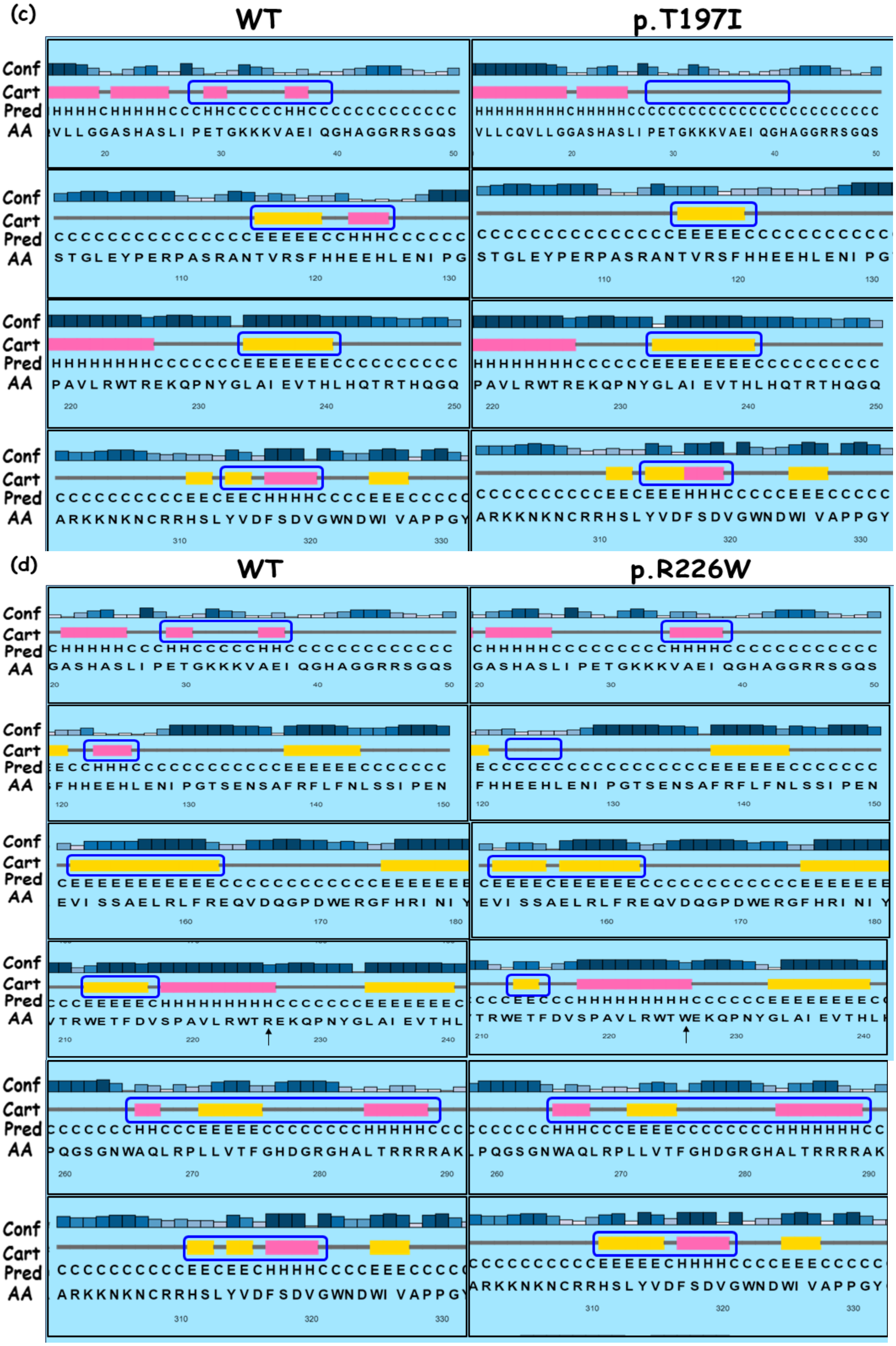
(c)
WT
p.T197I
(d)
WT
p.R226W
Conf
Cart
Pred
AA

**Supplementary Figure1. Comparative analysis of secondary structures of BMP4 wild-type (WT) and muteins (MUT) by Psipred server. (a-d) Predicted secondary structures of BMP4 WT and muteins (p.R113G, p.E151V, p.T197I, and p.R226W). The alterations in structures- helix (pink boxes) and strand (yellow boxes) has been masked by blue rectangles in both WT and MUTs.**

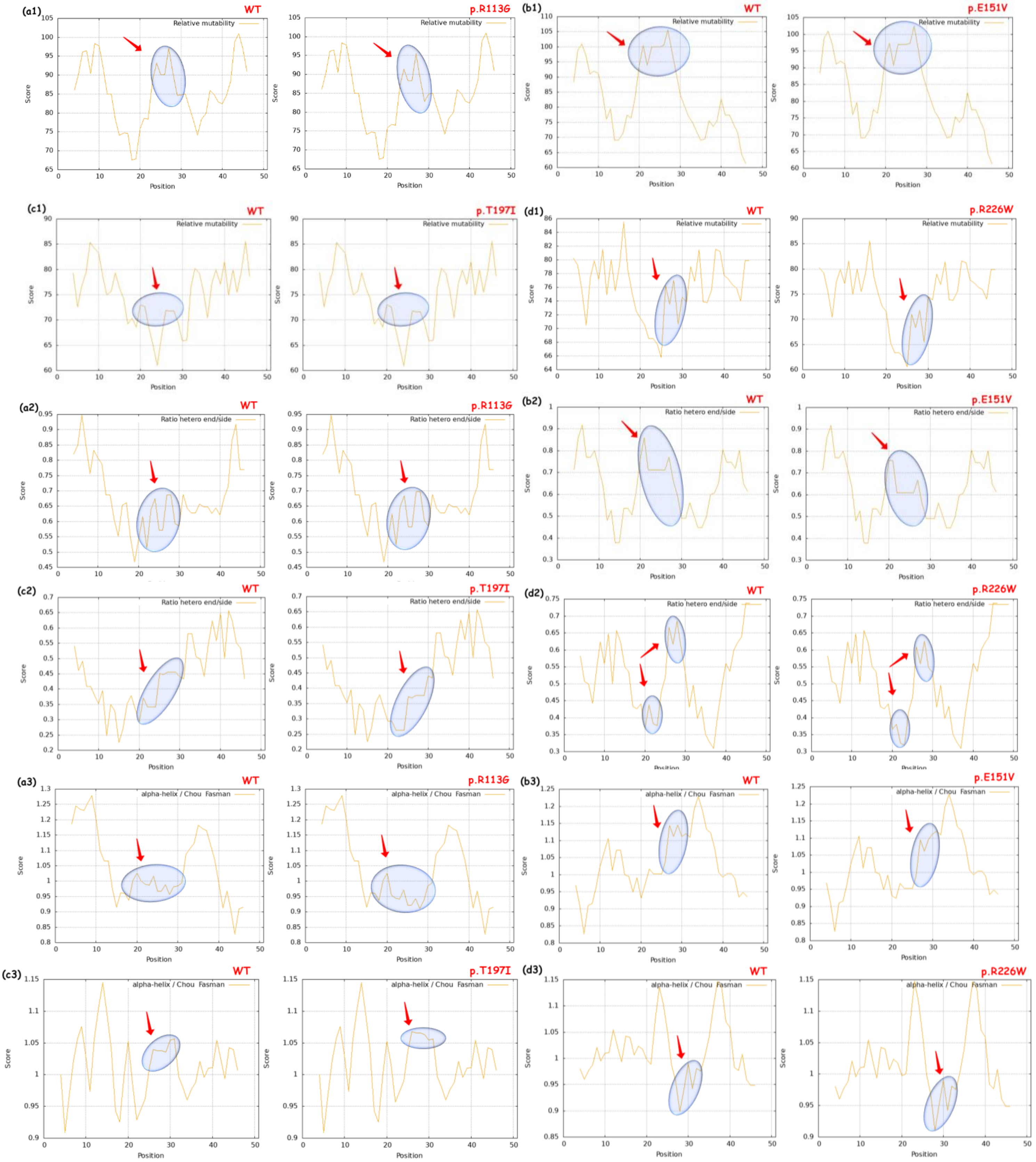

(a1)
WT
p.R113G
Relative mutability
(b1)
WT
p.E151V
Relative mutability
(c1)
WT
p.T197I
Relative mutability
(d1)
WT
p.R226W
Relative mutability
(a2)
WT
p.R113G
Ratio hetero end/side
(b2)
WT
p.E151V
Ratio hetero end/side
(c2)
WT
p.T197I
Ratio hetero end/side
(d2)
WT
p.R226W
Ratio hetero end/side
(a3)
WT
p.R113G
alpha-helix / Chou Fasman
(b3)
WT
p.E151V
alpha-helix / Chou Fasman
(c3)
WT
p.T197I
alpha-helix / Chou Fasman
(d3)
WT
p.R226W
alpha-helix / Chou Fasman
Score
Position

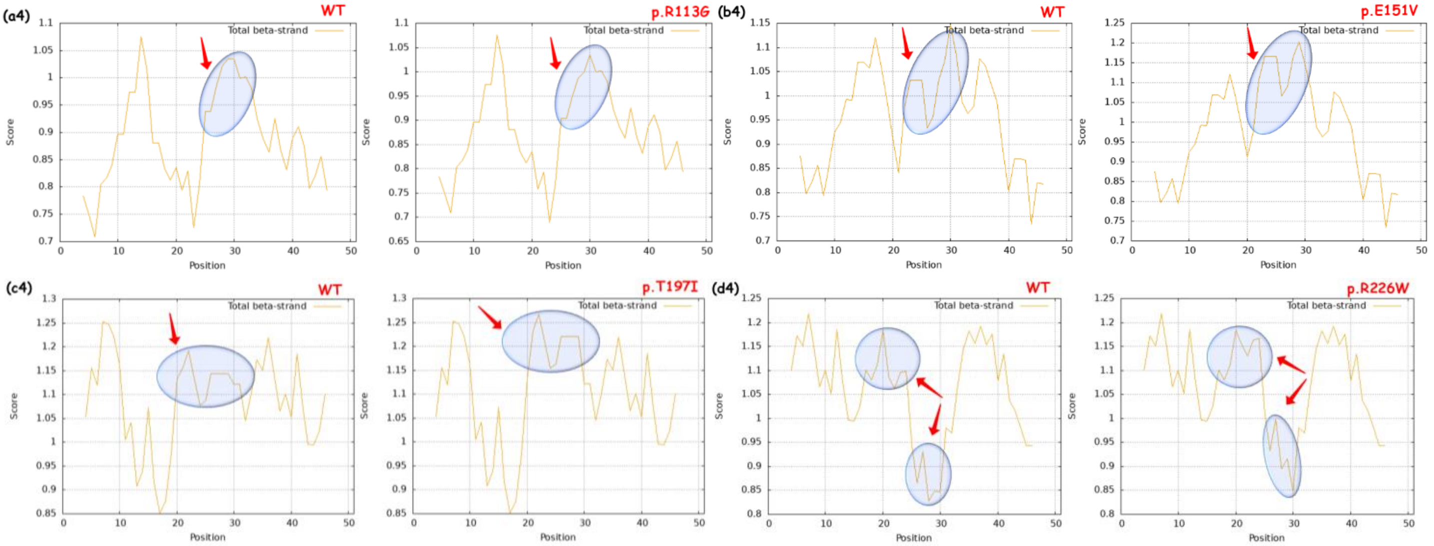


**Supplementary Figure2. Computational analysis of different physio-chemical properties of BMP4 WT vs muteins. (a1-d1) showing tendencies of relative mutability of WT and all the four muteins, (a2-d2) representing change in ratio hetero end side of muteins over WT, (a3-d3) comparative analysis of alpha-helix, (a4-d4) prediction of total -beta strand, of WT and muteins. The encircled regions marked with arrow representing the changes. WT= wild-type.**
